# The role of ecology and geography in the evolution of habitat isolation and sexual isolation among sister species of host-plant-specific insects

**DOI:** 10.1101/2022.01.02.474698

**Authors:** Linyi Zhang, Glen Ray Hood, James R. Ott, Scott P. Egan

## Abstract

While ecology and geography can play important roles in the evolution of reproductive isolation across the speciation continuum, the few studies to date examining their relative contributions have largely focused on the early stages of speciation. Studies exploring the evolution of multiple reproductive barriers among divergent lineages, where speciation is nearly complete, are also needed to gain a fuller understanding of the mechanisms promoting and constraining the speciation process. We examine the later stage of speciation by comparing the strength of habitat isolation and sexual isolation among closely related species of gall-forming wasps in the genus *Belonocnema* experiencing divergent ecologies due to different hos plant use and variable rates of migration due to geography. We found that the strength of both habitat isolation and sexual isolation is lowest among allopatric species pairs with the same host plant association, followed by allopatric species with different host plant association, and highest between sympatric species with different host-plant associations. This pattern strongly suggests that both ecology due to divergent host use, and geography contribute to the evolution of these two reproductive barriers. Notably, reproductive character displacement contributes to nearly half of the strength of both habitat isolation and sexual isolation in sympatry.

## Introduction

A central issue in speciation research is to understand the evolutionary forces that are responsible for generating reproductive barriers between diverging lineages (Wright 1978; Coyne and Orr 2004; Nosil 2012). Divergent selection driven by differences in ecology has been proposed to play an important role in promoting reproductive isolation (RI), a process termed as ecological speciation (Schluter 2000; Rundle and Nosil 2005). For example, phenological change driven by adaptation to different host plant species can lead to temporal isolation and thus reduced probability of mating between lineages of insect herbivores (Craig et al. 1993; Feder et al. 1993; Dopman et al. 2010; Hood et al. 2019; Zhang et al. 2019). Thus far, the majority of studies of ecological speciation have focused on the early stages of speciation, where partial reproductive barriers arise between populations from different environments (Funk et al. 2006; Nosil 2007; Kulmuni et al. 2020). However, it is less clear whether, and to what extent, ecology contributes to the later stages of speciation and the completion of RI in most systems.

Ecology may complete the speciation process by interacting with geography (Nosil 2007, 2012; Schwartz et al. 2010; Doellman et al. 2020). When lineages are in geographic contact with the potential for gene flow (e.g., sympatric or parapatric), selection can favor increased prezygotic isolation (Nosil 2007; Butlin and Smadja 2018) if there is a fitness cost to hybridization (e.g., extrinsic hybrid inviability), a process known as reinforcement (Servedio and Noor 2003). In addition, direct selection against immigrants from alternative environments can also promote increased habitat isolation by favoring individuals with high fidelity for their native environment (Nosil et al. 2006). These two mechanisms can occur in lineages experiencing migration, and under these conditions, sympatric and/or parapatric lineages are predicted to exhibit higher prezygotic isolation than allopatric lineages with little or no gene flow, a pattern known as reproductive character displacement (Brown and Wilson 1956; Butlin 1987a; Noor 2001). Alternatively, in the absence of divergent selection, gene flow in sympatry can erode RI by homogenizing lineages (Nosil et al. 2003). Despite the importance of geography in promoting or constraining speciation (Rosser et al. 2019; Jiménez-López et al. 2023), many studies that have investigated ecological speciation have done so without specifically addressing and isolating its role (e.g., Via et al. 2000; Nosil et al. 2002; Lowry et al. 2008b; Egan et al. 2012a). Moreover, there is growing consensus that the completion of speciation commonly requires the evolution of multiple reproductive barriers that cumulatively reduce gene flow to near zero between diverging lineages (Ramsey et al. 2003; Nosil 2007; Dopman et al. 2010; Richards and Ortiz-Barrientos 2016; Lackey and Boughman 2017; Zhang et al. 2021b). Thus, to examine the potential effect of ecology and geography on the completion of speciation, multiple reproductive barriers between closely related, but ecologically divergent taxa experiencing variable levels of migration due to geographic context must be investigated (e.g., *Timema* stick insects Nosil 2007).

A powerful approach to evaluating the absolute and relative contributions of ecology and geography to the completion of speciation is to compare the strength of RI between lineages characterized by different combinations of ecology and geography, as illustrated in Fig. 1 (Nosil 2007). For scenario one, measurable RI detected between allopatric lineages that experience the same ecological conditions suggests the role of genetic drift via geographic isolation, other sources of selection not directly associated with host plant use, and different evolutionary histories (Dobzhansky 1934; Funk and Nosil 2008). Importantly, the levels of RI observed for this scenario serve as the baseline for comparisons of RI under the two alternative scenarios. For scenario two, if allopatric lineages from different environments are observed to exhibit higher RI than allopatric lineages from the same environment, the inference is that divergent selection has acted over and above other evolutionary mechanisms to promote RI when gene flow is low or not present. Lastly, for scenario three, reproductive character displacement, where sympatric lineages from different environments exhibit elevated RI compared to allopatric lineages from different environments, is consistent with the inference that selection interacts with gene flow (i.e., reinforcement) to promote increased prezygotic RI (Noor 1999). Alternatively, the observation of lower RI among lineages from different environments in sympatry compared to allopatry supports the inference that gene flow constrains the evolution of RI (Nosil et al. 2003). Few studies have applied this specific approach despite the potentially important mechanistic insights into the evolution of RI provided by this comparative framework (Nosil 2007).

**Figure 1.**
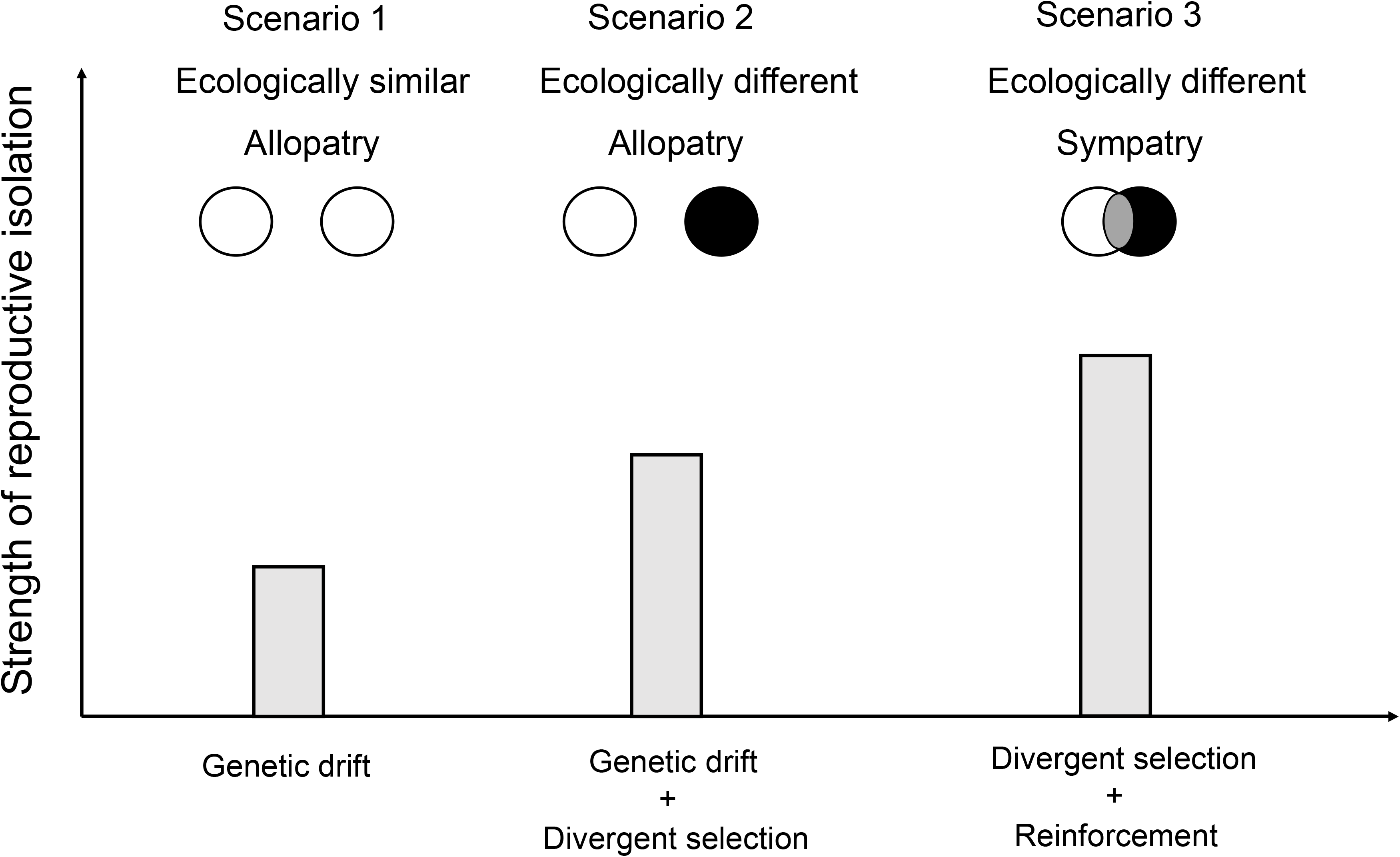
Illustration of possible evolutionary mechanisms that promote reproductive isolation (RI) across different modes of speciation (modified from Nosil 2007). Scenario 1 illustrates the strength of RI between two allopatric lineages that experience ecologically similar environmental conditions. Possible evolutionary mechanisms in promoting RI in this scenario including genetic drift via geographic isolation, other sources of selection not directly associated with divergent ecology, and different evolutionary histories. Importantly, RI observed in Scenario 1 serve as the baseline for comparisons of RI under the two alternative scenarios. Scenario 2 illustrates the strength of RI between two allopatric lineages that experience divergent selection from differing ecological environments plus all the phenomena capture in scenario 1. Scenario 3 illustrates the strength of RI between two sympatric lineages that experience divergent selection from differing ecological environments, where potential opportunities for gene exchanges might promote or hamper the evolution of reproductive isolation. The higher values of RI in Scenario 2 compared to Scenario 1 suggest that divergent selection promotes RI. Higher values of RI in Scenario 3 compared to Scenario 2 suggest that selection that occurs in sympatry, such as reinforcement, elevates RI. Alternatively, the observation of lower RI among lineages from different environments in sympatry compared to allopatry supports the inference that gene flow constrains the evolution of RI.

Herein, we apply the comparative framework illustrated in Fig. 1 to determine the roles of ecology and geography by comparing the strength of both habitat isolation and sexual isolation among a complex of three host-plant-specific species of gall-forming wasps in the genus *Belonocnema* [Hymenoptera: Cynipidae]. These three species are each locally adapted to live oak species in the genus *Quercus* (subsection *Virentes*). Two species, *B. treatae* and *B. fossoria*, co-occur over a large portion of their geographic ranges in the coastal southeastern United States on their respective host plants, the southern live oak, *Q virginiana* (*Qv*), and the sand live oak, *Q. geminata* (*Qg*) (Fig. 2) (Driscoe et al. 2019; Zhang et al. 2021c). In the western-most range of *Qv* along the Gulf coast, a third species, *B. kinseyi*, also develops on *Qv* where it is allopatric to both *B. treatae* and *B. fossoria*. Thus, both the geographic distribution of host plants and the patterns of host plant use by *Belonocnema* (allopatric vs. sympatric lineages, and ecologically similar vs. ecologically different lineages), allow us to apply this comparative framework (Nosil 2007) to explore the evolutionary mechanisms underlying two prezygotic reproductive barriers simultaneously. Specifically, we assess the strength of RI between the following species pairs of *Belonocnema* to evaluate patterns of RI as a function of differing environments and geographic context: *B. kinseyi* × *B. treatae* (allopatric species pair with the same ecology), *B. kinseyi* × *B. fossoria* (allopatric species pair with different ecology), and *B. treatae* × *B. fossoria* (sympatric species pair with different ecology).

**Figure 2.**
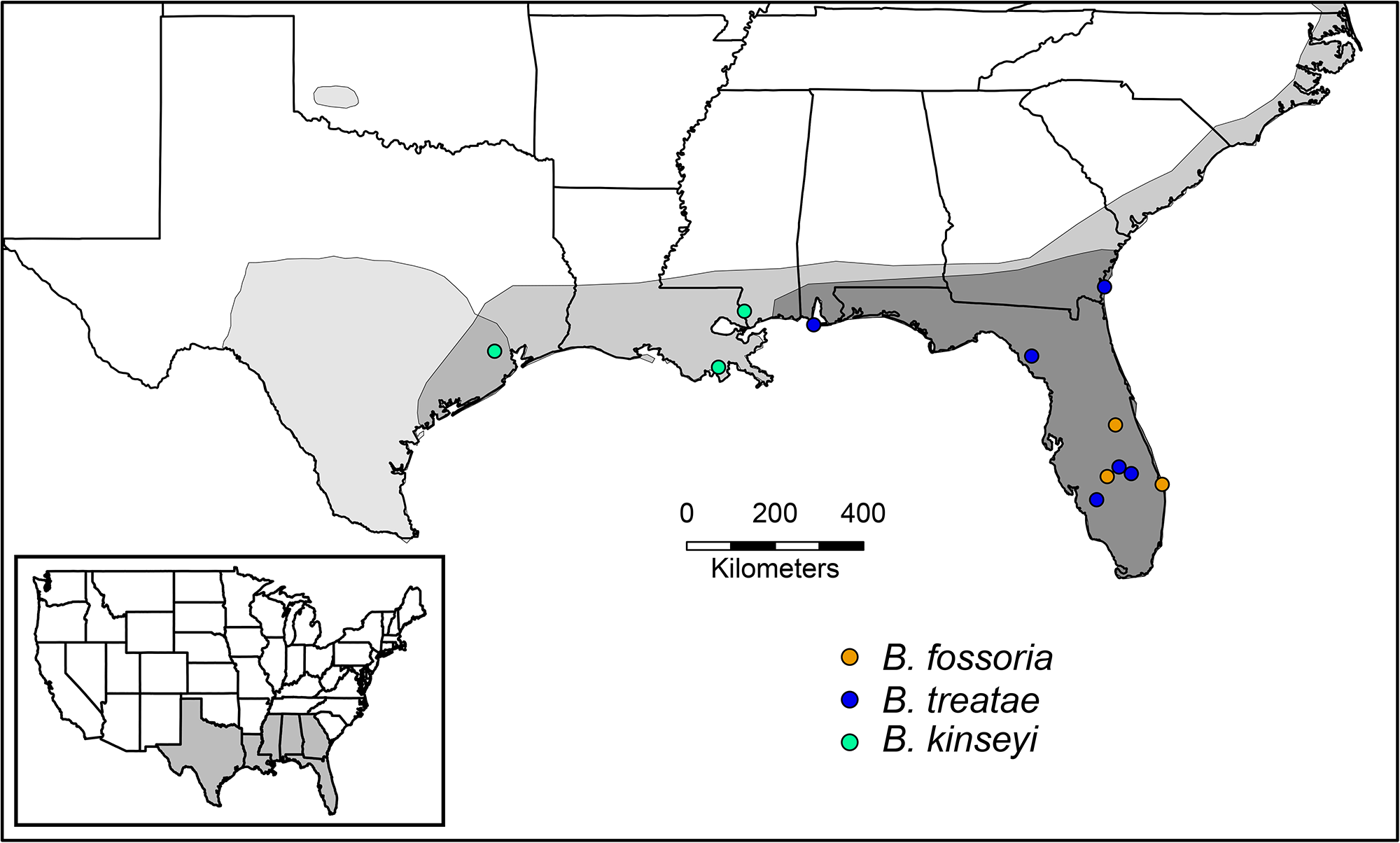
The geographic ranges of host plants used by gall wasps in the genus *Belonocnema*: *Q. fusiformis* (restricted mainly to central and south Texas; light grey)*, Q. virginiana* (distributed broadly across the coastal southeastern U.S.; intermediate grey), and *Q. geminata* (restricted mainly to Florida; dark grey) (modified from Driscoe et al. 2019). Also shown are the collection sites where different species of sexual generation adult *Belonocnmea* were sampled for habitat isolation and sexual isolation experiments: green, (*B. kinseyi*) blue, (*B. treatae*) and orange (*B. fossoria*). This color scheme is used throughout all Figures to represent each *Belonocnema* lineage.

## Methods

### Study system

The genus *Belonocnema* (Hymenoptera: Cynipini: Cynipidae) comprises three gall wasp species that specialize on American live oaks of the southern United States (genus *Quercus*; subsection Virentes) (Zhang et al. 2021c). As with many cynipids, these species exhibit cyclical parthenogenesis whereby the sexual and asexual generations alternate seasonally to complete an annual life cycle (Lund et al. 1998). In the spring, sexual generation adult males and females developing separately in single sex, multi-chambered root galls emerge, mate immediately and oviposit into newly flushed leaves. Oviposition at the onset of new leaf growth is critical for successful asexual generation gall initiation (Zhang et al. 2017; Hood et al. 2019). Successful oviposition and larval establishment results in a single-chambered, spherical gall that forms on the underside of leaves. In early fall, asexual adults emerge from leaf galls within the canopies of trees and descend to the ground and search for root tissue suitable for gall induction (Hood and Ott 2010). This study focuses on estimating RI among sexual generation individuals, where habitat isolation and sexual isolation can directly affect gene flow among the three *Belonocnema* species.

The geographic ranges of the two host plant species, *Q. virginiana* and *Q. geminata*, broadly overlap in the southeastern U.S., however the geographic range of *Qv* also extends westward along the Gulf coast to central Texas (Fig. 2). Despite general geographic sympatry, *Qv* and *Qg* occupy different microhabitats, where *Qg* occurs in drier, higher pH, sandy-like soil compared to *Qv* that occurs in moist, lower pH soil. In addition, *Qv* and *Qg* also differ in the phenology of spring leaf flush, physiological traits, tree height and leaf morphology (Cavender Bares and Pahlich 2009; Cavender Bares et al. 2015).

Four additional lines of evidence, in addition to the geographic distribution and host-association described above, establish the suitability of the study system for comparative analysis of RI among the species pairs to isolate the role of ecology and geography during the late stages of speciation. First, the sympatric species pair *B. treatae* and *B. fosoria* exhibit near complete RI (range among populations = 0.95-0.99) contributed to by multiple reproductive barriers, including temporal isolation (Hood et al. 2019), sexual isolation (Egan et al. 2012a), and habitat isolation (Egan et al. 2012b). As a result, high genomic divergence between the two species at multiple markers is found across the genome (G_st_ = 0.37; Driscoe et al. 2019). Second, the two different host plant species have been documented to exert strong divergent selection on the *Belonocnema* species, as supported by the observation that both immigrant and hybrid gall forming wasps suffer significant reductions in fitness dependent on the host plant species (Zhang et al. 2017, 2021a,b; Fig. S1). Third, population genomic analysis supports the expectation that migration can occur between the sympatric species (Driscoe et al. 2019). Lastly, allopatric *B*. *kinseyi* are more genetically distant from both sympatric *B. treatae* and *B. fossoria*, as inferred by both mtDNA and population genomic data (Egan et al. 2012a, Driscoe et al. 2019).

### Sample collection

In the spring of 2016 through 2019, we collected mature root galls containing pupal stage individuals of each gall former species from their respective host plants: *B. fossoria* (3 sites), *B. treatae* (6 sites), and *B. kinseyi* (3 sites) (Fig. 2; Supplemental Table 1). To secure unmated females, each root gall was individually placed in a 30 ml clear plastic vial and stored at room temperature. The timing of peak emergence of *B. treatae* occurs two to three weeks earlier than *B. fossoria* (Hood et al. 2019). Therefore, to synchronize emergence for experimental purposes, we placed one-half of the root galls collected from *Qv* at each site into a refrigerator at 4°C for one week to delay the emergence of *B. treatae*. The vials were monitored every two days and upon emergence, one-to two-day-old adults were collected and used to perform the host preference and mate choice tests described below. All tests were conducted under controlled greenhouse conditions at Rice University, Houston, Texas, in 2016, 2018, and 2019, and in ambient conditions in a screened-in outdoor enclosure at Archbold Biological Station, Venus, Florida, in 2017.

### Estimating habitat isolation

To evaluate the relative roles of ecology (different host plant associations) and differences in geography (different opportunities for migration and gene flow) on habitat isolation, we measured host plant preference for the two sympatric species, *B. treatae* and *B. fosoria*, as well as for the allopatric species, *B. kinseyi*, using a controlled two-choice experimental approach following Egan et al. (2012*b*) (Table S1). As female host choice directly determines the environment in which oviposition and larval development proceeds for the next generation, and male *Belonocnema* display lower host fidelity (Egan et al. 2012b), we only used females to quantify host preference. In 2016 and 2017, host preference trials were conducted within sealed 500 mL clear-plastic cups that contained a fresh 15–20 cm stem cutting of both *Qv* and *Qg* from Florida with a similar number and size of leaves to minimize potential effects of differences in plant biomass on host choice. In 2019, trials were conducted within 60 × 15 mm (diameter × height) Petri dishes stocked with a single newly flushed leaf of equivalent size of each host plant positioned at opposing sides. For all trials in each year, a single unmated female was aspirated into each container and then observed at 2-minute intervals for 30 minutes for a total of 15 observations. At each interval, we recorded the location of the female as being on *Qv*, on *Qg,* or on the surface of the cup or Petri dish. In total, 557 host preference assays were conducted using 126 *B. treatae* and 130 *B. fossoria* from sympatric *Qv* and *Q*g sites and 301 allopatric *B. kinseyi* from *Qv* sites. Only those trials in which individuals were observed on a host plant at least once were retained for analysis (76.5% of trials, Table S1). In total, we made 8,355 individual observations over 278.5 testing hours for the assessment of host preference.

To compare habitat isolation among species, we conducted two parallel analyses. First, we calculated host preference of each individual by dividing the number of time points where a female was observed on its natal host plant by the total number of time points observed on either host plant during each trial. These values of host preference range from 0 to 1, with values less than 0.5 indicating increasing preference for the non-natal host plant, values of 0.5 indicating equivalent preference for both natal and non-natal host plants, and values greater than 0.5 indicating preference for the natal host plant. The host preference value of each species was then compared using a generalized linear mixed model (GLMM) with the response variable ‘host preference’ assigned a beta-binomial distribution to control for overdispersion of the binomially distributed data (Kim and Lee 2017; Lenth et al. 2020). The independent variables ‘arena type’ (plastic cup or Petri dish) and ‘gall wasp species’ were included as fixed effects, with ‘collection site’ treated as a random effect. This analysis allows us to explore the species level differences while ruling out population level variation.

Second, we used the above values of host preference expressed by each species to calculate and compare the strength of habitat isolation between species. Following the approach described in Nosil et al. (2005), the strength of habitat isolation (HI) was quantified as the absolute value of the difference in mean host preference towards *Qv* between each species. This species-pair metric ranges from 0 to 1, with 0 indicating similar habitat preferences and no habitat isolation and 1 indicating divergent habitat preferences and complete habitat isolation between species. For the purposes of statistical analysis, we obtained 10,000 bootstrap values of this metric of habitat isolation for each of the three species pairs, *B. kinseyi* × *B. treatae*, *B. kinseyi* × *B. fossoria*, and *B. treatae* × *B. fossoria*, by randomly resampling individual host preference values from the original data and recalculating habitat isolation (John and Fuller 2021). Bootstrap values of habitat isolation for the three heterospecific species pairs were then compared via t-tests. The significance level for multiple comparisons was adjusted to α = 0.05/3 = 0.016 using a Bonferroni correction (Haynes 2007).

### Estimating sexual isolation

To evaluate the relative roles of ecology (host plant association) and geography (opportunity for migration and gene flow) on sexual isolation, we measured sexual isolation for the two sympatric species (*B. treatae* and *B. fosoria*) as well as for the allopatric species, *B. kinseyi*, using no-choice preference trials (Table S3). Similar to the host preference tests, trials were conducted in 2016, 2017, and 2018 within 60 × 15 mm Petri dishes lined at the base with damp filter paper. For each trial, one male and one female were aspirated into the Petri dish and observed at 2-minute intervals for 30 minutes for a total of 15 observations. At each interval, we recorded three courtship and mating-related interactions: (a) whether the male was engaged in wing buzzing (or fanning), an important male courtship display in insects (Villagra et al. 2011), (b) whether the male mounted the female, a behavior that precedes copulation, and (c) whether the pair was copulating, defined as contact of male and female abdomens. In total, 1,123 mating assays were conducted using 225 female and 153 male *B. treatae* from *Qv*, 563 female and 455 male *B. fossoria* from *Q*g, and 335 female and 515 male *B. kinseyi* from *Qv* (see Table S3 for pairings and replication). In total, we recorded 16,845 observations over 561.5 testing hours for the assessment of mate preference.

Similar to the habitat isolation analysis, we used two complementary methods to compare sexual isolation among species pairs. First, we calculated and then tested whether the probability of mating differed between conspecific and heterospecific pairs for each set of species with larger differences in the probability of mating equating to greater sexual isolation between species pairs. The probability of mating for each individual pair was quantified as 0 if copulation did not occur during the observations and 1 if copulation occurred. Further dissection of male-specific (wing buzzing) and female-specific (allowing male to mount) mate preferences are included in the Supplemental analysis. The mating probability of the same species vs. different species treatments for each species pair was then compared using GLMM where the response variable ‘mating probability’ was considered to be binomially distributed, with the independent variables ‘collection year’ (2016, 2017, 2018 or 2019) and cross type (conspecific or heterospecific) included as fixed effects, and ‘collection site’ of males and females treated as a random effect. The three different heterospecific species pairs, *B*. *kinseyi* × *B. treatae*, *B. kinseyi B. fossoria*, and *B. treatae* × *B. fossoria*, were compared with the appropriate two conspecific mating trials for each species pairing (e.g., *B. kinseyi* × *B. kinseyi, B. treatae* × *B. treatae,* or *B. fossoria* × *B. fossoria*). Due to the lack of independence, the interaction between species pair type and individual pair cannot be directly tested using GLMM. Thus, while GLMM analysis allows us to isolate the collection sites effects on mating probability, we also quantified and compared differences in the strength of sexual isolation (SI) between species pairs using the method described by Sobel and Chen (2014): SI = 1 – 2 × [heterospecific mate frequency / (heterospecific mate frequency + conspecific mate frequency)]. In this metric, a value of zero indicates no sexual isolation whereas a value of 1 indicates complete sexual isolation. Similar to the analysis of habitat isolation, we then generated 10,000 bootstrap values of sexual isolation between each of the three species pairs by randomly resampling mated and unmated pairs from the original data and recalculating sexual isolation. The bootstrap values of sexual isolation for the three heterospecific species pairs were then compared via t-tests with the significance level for the multiple comparison adjusted to α = 0.05/3 = 0.016 using a Bonferroni correction.

All GLMM analyses in this study were followed by multiple pairwise comparisons among *Belonocnema* lineages with different geographic and host plant associations using a Tukey’s post hoc test with the function *lsmeans* in package ‘emmeans’. All analyses including bootstrapped simulations were conducted in R version 4.0.2 (R core team 2021).

### Relative contributions of ecology and geography to habitat isolation and sexual isolation

Following the framework outlined in Fig. 1, the contributions of different evolutionary mechanisms to habitat isolation and sexual isolation were assessed across different species pairs. The contribution of genetic drift via geographic isolation or other mechanisms such as selection not directly associated with host plant use, and/or different evolutionary histories (Dobzhansky 1934; Funk and Nosil 2008) was estimated using allopatric species with the same host plant association (*B. kinseyi* and *B. treatae*, Fig. 1, scenario one). The contribution of ecology highlighted in scenario two (Fig. 1) was estimated as the difference in RI between the allopatric species pair with different hosts (*B. kinseyi* × *B. fossoria*) compared to the allopatric species pair on the same host (*B. kinseyi* × *B. treatae*) (i.e., the difference between scenario 1 and scenario two in Fig.1; similar to Funk 1998, Nosil et al. 2002). Lastly, the contribution of mechanisms acting in sympatry (e.g., reinforcement) was estimated as the difference in RI between the sympatric species pair on different hosts (*B. treatae* × *B. fossoria*) versus the allopatric species pair on different hosts (*B. kinseyi* × *B. fossoria*) (i.e., the difference between scenario three and scenario two; Fig. 1). The values of habitat isolation (HI) and sexual isolation (SI) used in these comparisons are taken as the mean of 10,000 bootstrap values obtained from the previous analyses described above. To convert these absolute values to their relative contributions, each value was divided by the total RI for that barrier.

## Results

### Strength of habitat isolation

Populations of *B. treatae* developing on *Qv* that are sympatric with *B. fossoria* displayed significantly higher host plant fidelity (mean ± SE, 0.778 ± 0.042) than allopatric populations of *B. kinseyi* developing on *Qv* (0.647 ± 0.023, *t* = 2.49, *P* = 0.035, Fig. 3A) and sympatric populations of *B. fossoria,* specialized on *Qg* (0.626 ± 0.037, *t* = 2.56, *P* = 0.029, Fig. 3A). Interestingly, the individual trials performed in the smaller Petri dishes exhibited significantly higher host preference than the larger cups (*P* < 0.01) suggesting proximity to the two host plant options increased the expression of host preference. However, testing environment did not affect the overall difference in host preference among different *Belonocnema* species, as indicated by the interaction term in the GLMM (*P* = 0.356, Table S4), suggesting that this difference is unlikely to affect our qualitative interpretation of the comparisons between species. Correspondingly, we found the strength of habitat isolation to be the lowest between allopatric populations of *B. kinseyi* and *B. treatae* that share the same host plant species (mean ± 95% CI: 0.157, 0.064–0.248), followed by allopatric populations of *B. kinseyi* and *B. fossoria* (0.183, 0.099–0.265) that use different host plant species, and sympatric populations of *B. treatae* and *B. fossoria* (0.340, 0.235–0.442) that use different host plant species (Fig. 3B).

### Strength of sexual isolation

The three *Belonocnema* species exhibited significantly higher probability of mating with conspecifics compared to heterospecifics that used different host plants (*B. kinseyi* × *B. fossoria*: *Z* = 2.511, *P* = 0.012; *B. treatae* × *B. fossoria*: *Z* = 2.279, *P* = 0.023; Fig. 4A). In contrast, the probability of mating did not differ among allopatric conspecific and heterospecific *B. kinseyi* and *B. treatae*, which share the same host plant species (*Z* = 0.877, *P* = 0.380; Fig. 4A). Correspondingly, the strength of sexual isolation was lowest between allopatric *B. kinseyi* and *B. treatae* using the same host plant species (mean ± 95% CI: 0.081, -0.137–0.302), but was ∼1.7 times greater between allopatric *B. kinseyi* and *B. fossoria* using different host plant species (0.138, 0.013–0.259). Lastly, the strength of sexual isolation was highest between sympatric populations of *B. treatae* and *B. fossoria* using different host plants (0.337, 0.068–0.598). Further dissection of male and female specific mate preferences, which are similar to the general patterns described here, are included in the Supplement (Table S5, Table S6, and Fig. S2).

**Figure 3.**
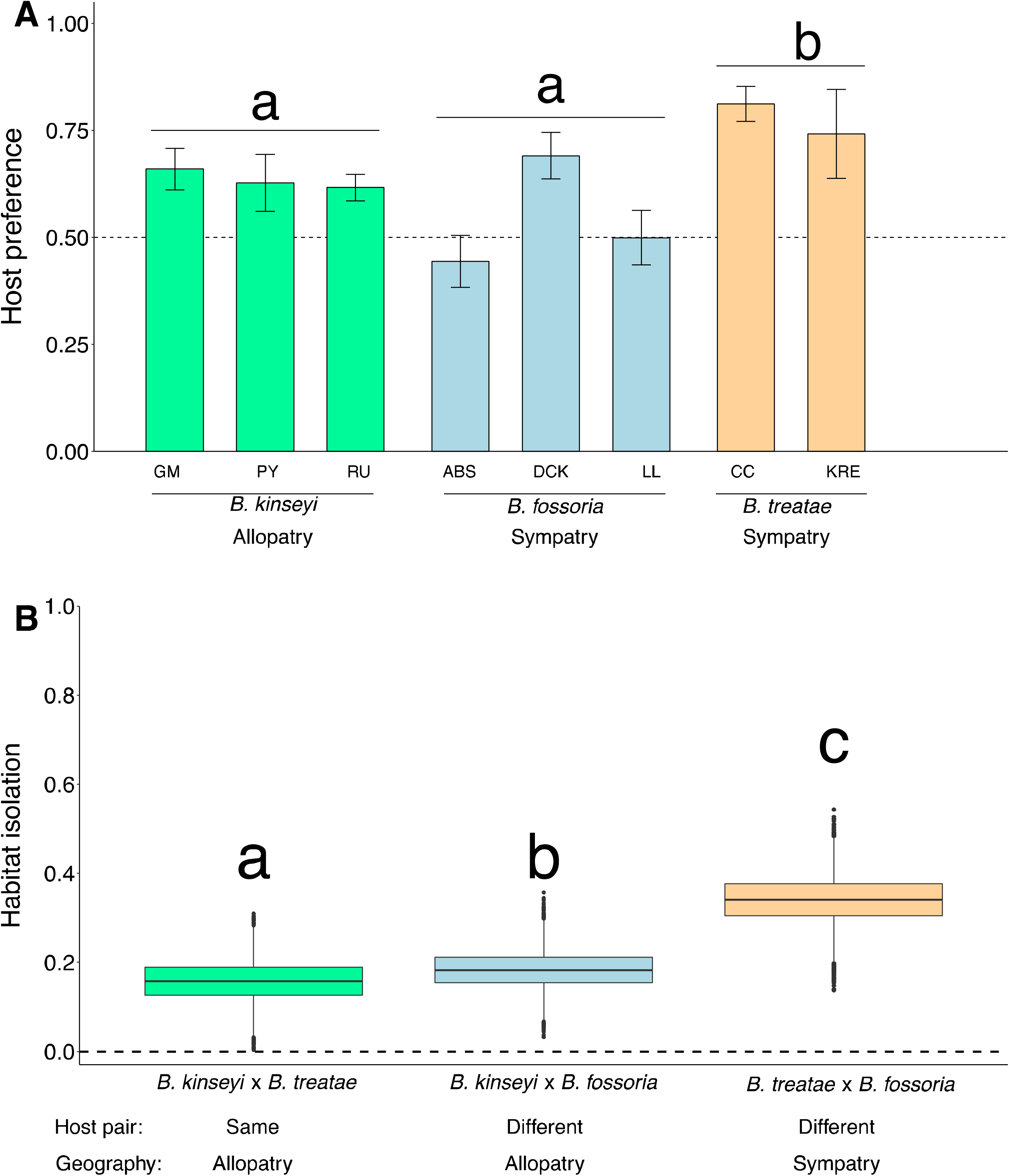
**(A)** Host preference (mean ± SE) defined as the percent of time females spent on their native host plant when given the choice of native and novel host plants evaluated for *B. kinseyi*, *B. fossoria*, and *B. treatae* populations (see Fig. 2 and Table S2 for population identification details). The horizontal dashed line at 0.5 represents the null hypothesis of no preference for either host plant. Letters above bars indicate statistically significant differences among species (*P* < 0.05). **(B)** Boxplot of 10,000 bootstrap values of habitat isolation for each of three comparisons between pairs of species, *B. kinseyi* × *B. treatae*, *B. kinseyi* × *B. fossoria*, and *B. treatae* × *B. fossoria*, oriented along the x-axis in the same way as our conceptual framework in Figure 1. The horizontal dashed line at 0 represents the null hypothesis of no habitat isolation. Different letters above boxplots indicate statistically significant difference among the three species pairs (*P* < 0.05).

**Figure 4.**
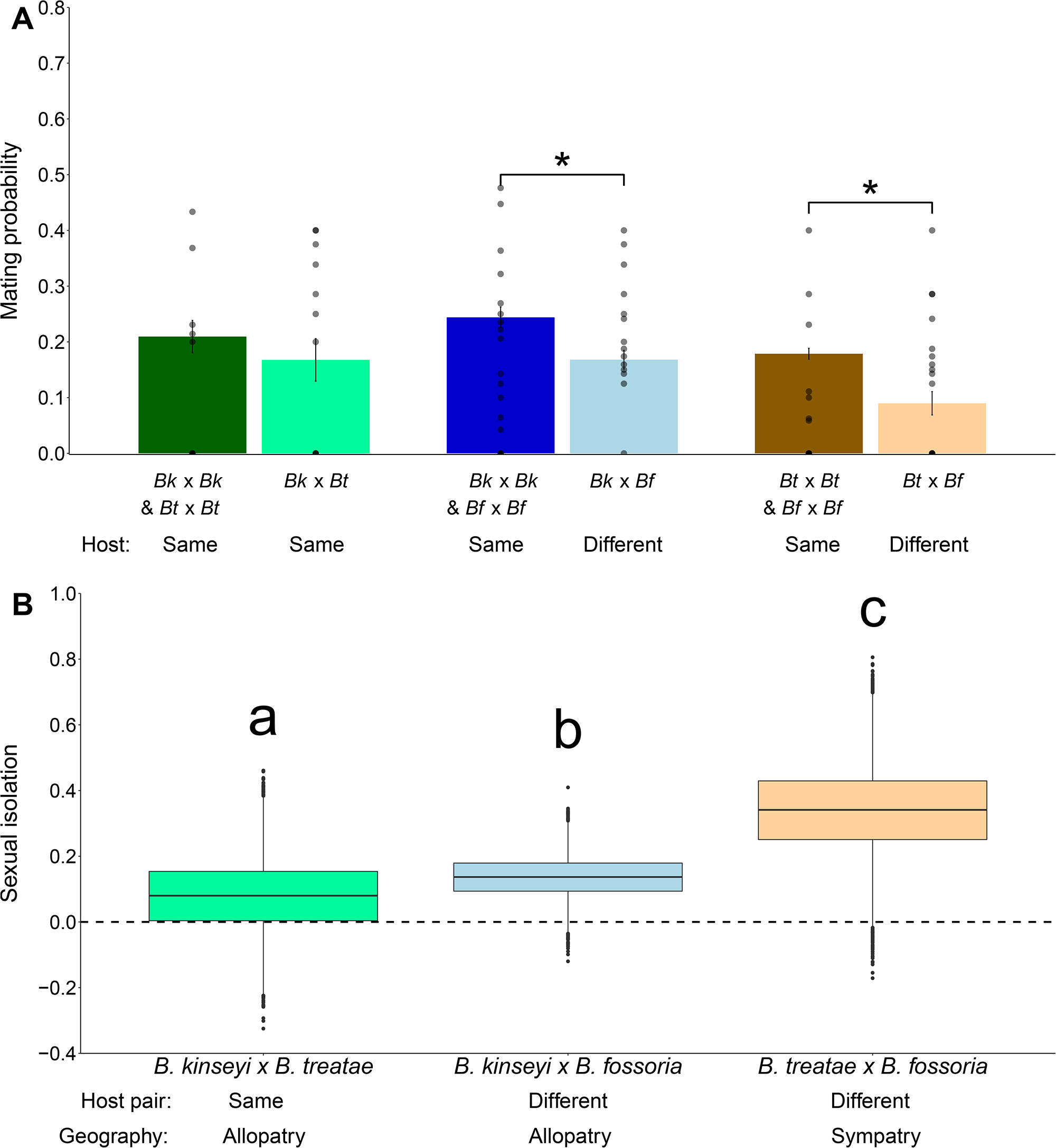
**(A)** Boxplot of the probability of mating among wasp species in the genus *Belonocnema* that use the same and different host plant species, as determined by sexual isolation experiments. The bar values are the least squared means of mating probability within the group with standard error evaluated from GLMMs. The dots represent the means of mating probability for each population pair. *Bk*, *Bt*, and *Bf* identify *B. kinseyi*, *B. treatae*, *B. fossoria*, respectively. **(B)** Boxplot of 10,000 bootstrapped values of the strength of sexual isolation for the three comparisons between species pairs, *B. kinseyi* × *B. treatae*, *B. kinseyi* × *B. fossoria*, and *B. treatae* × *B. fossoria*, oriented along the x-axis in the same way as our conceptual framework in Figure 1. The dashed line at 0 identifies the null hypothesis of no sexual isolation. Different letters above the boxplots indicate statistically significant difference between species pairs (*P* < 0.05).

### Relative contributions of ecology and geography to habitat isolation and sexual isolation

Using the comparative framework in Fig. 1 and the values of habitat isolation and sexual isolation documented in Fig. 3B and 4B for each type of species comparison (Scenario one: same-host, allopatric comparison; Scenario two: different-host, allopatric comparison; Scenario three: different-host, sympatric comparison), we calculated the relative contributions of three components to each overall barrier: (1.) Genetic drift and/or non-host-associated divergence, (2.) host-associated divergence, and (3.) character displacement in sympatry (e.g., reinforcement). For habitat isolation, the relative contributions of genetic drift and/or non-host-associated divergence was 46%, host-associated divergence was 8%, and character displacement in sympatry was estimated to be 46% of overall HI (Fig. 5). For sexual isolation, the relative contributions of genetic drift and/or non-host-associated divergence was 24%, host-associated divergence was 16%, and character displacement in sympatry was estimated to be 60% of overall SI (Fig. 5).

**Figure 5.**
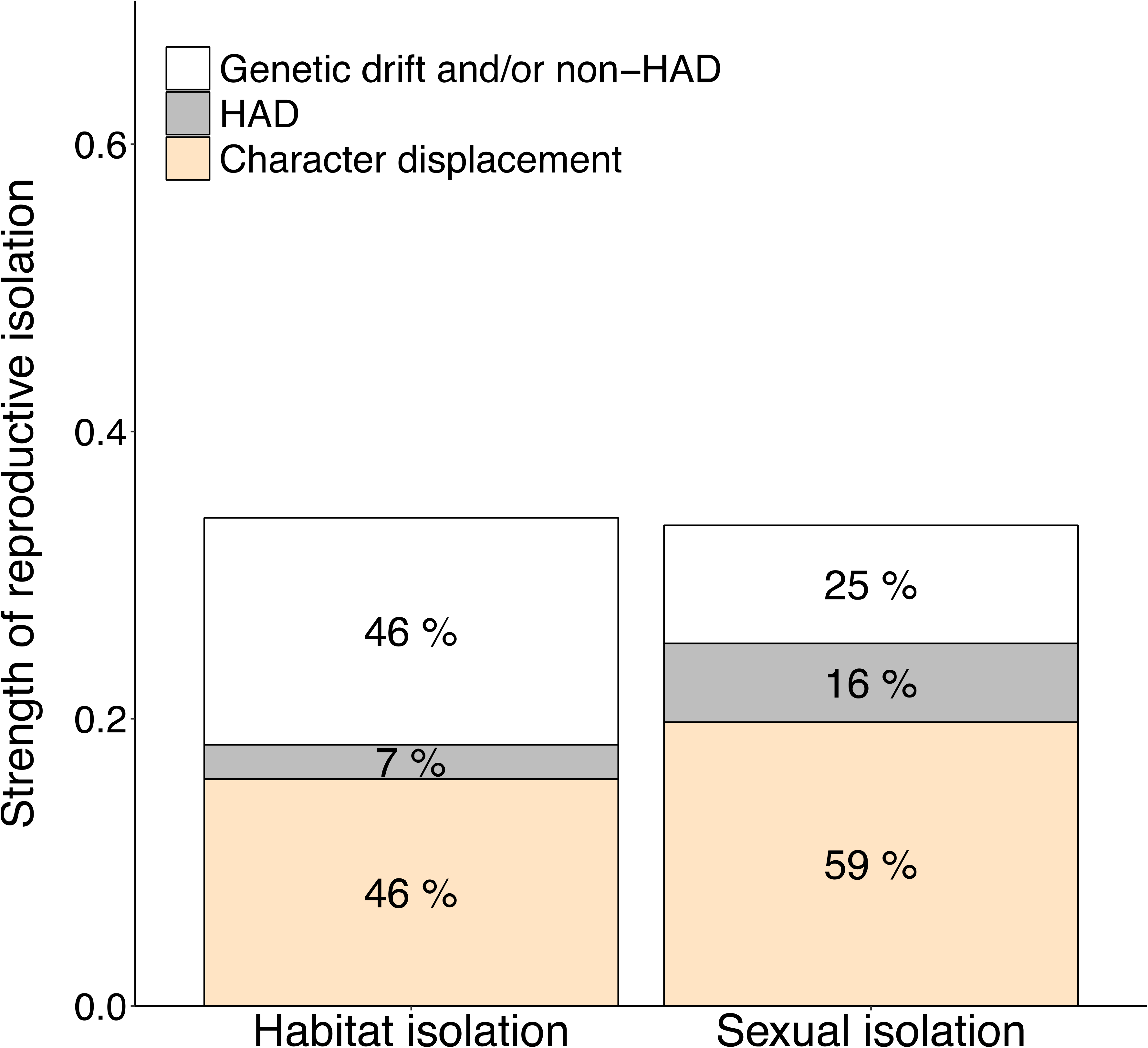
Estimates of the relative contribution of different evolutionary mechanisms promoting habitat isolation and sexual isolation across different species pairs: Genetic drift and/or non-HAD, HAD, and character displacement. (HAD = host-associated divergence) (see comparative framework in Figure 1 for contrasts calculated here).

## Discussion

In this study, we adopted a comparative framework (Fig. 1) to infer the separate and interactive contributions of ecology and geography to the evolution of habitat isolation and sexual isolation, as proposed by Nosil 2005, 2007. These two prezygotic barriers constitute important components of RI among the three closely related species of *Belonocnema* gall wasps (Egan et al. 2012a,b; Hood et al. 2019). Among the limited cases where the evolutionary mechanisms underlying RI have been explored in this context, the studied lineages display only partial RI, and are thus considered to be at an early stage of speciation (e.g., Nosil 2007). However, as suggested by Kulmuni et al. (2020), studies of more divergent taxa can add to our overall understanding of the speciation process by examining the evolutionary factors that may be important to completing speciation. In this study, we inferred how divergent selection due to host plant use (ecology) and selection against migration/hybridization in sympatry (geography) might influence the evolution of reproductive barriers among three closely related species of gall-forming insects and add an important and much needed case study of these evolutionary mechanisms at the later stages of speciation.

For both habitat and sexual isolation, we found that the strength of RI between species was lowest among allopatric species that shared the same host plant association, higher between allopatric species with different host plant associations, and highest between sympatric species with different host plant associations as illustrated in Fig. 3 and 4. This pattern of RI in relation to geographic context and host plant association corresponds to predictions of the conceptual model of Nosil (2007), which is predicated on direct and indirect selection processes. In addition, it leads to new inferences in this system regarding the relative roles of different mechanism at the later stages of speciation (Kulmuni et al. 2020).

Our finding of significant habitat isolation and sexual isolation among allopatric species that share the same host plant association suggests that neutral processes associated with genetic drift and/or selection that is not directly associated with host plant use can contribute to the evolution of both reproductive barriers (Figure 5). The unexpectedly high habitat isolation and sexual isolation between allopatric species with the same host plant association (46% of total HI, 24% of total SI) indicates a potentially more complicated evolutionary story in a system where divergent selection from different host plants has been the main focus (e.g., Egan et al. 2021a, 2012b, Egan et al. 2013, Zhang et al. 2017, Hood et al. 2019, Driscoe et al. 2019, Zhang et al. 2021a, 2021b). Such high prezogytic isolation between allopatric speices pairs could be a result of neutral process such as genetic drift, or from selection not directly associated with host plant use. For instance, reproductive isolation could be an indirect product of the fixation of different advantageous mutations in separate populations experiencing similar selection, a process known as mutation-order speciation (Schluter 2009; Nosil 2012). In the case of allopatric *B. treatae* and *B. kinyesi* that share the same host plant, each lineage may have developed different phenotypic pathways underlying adaptation to the same host plant environment, and these differing solutions could generate assortative mating within lineages leading to partial habitat and/or sexual isolation (Schluter 2009; Nosil and Flaxman 2011). Given the complex genetic pathways involved in gall formation, it might not be surprising that lineages differ in their adaptations to manipulate their host (Martinson et al. 2022).

The higher estimates of habitat isolation and sexual isolation observed between allopatric species feeding on different host plants compared to allopatric species feeding on the same host plant species strongly suggest that divergent selection can promote RI regardless of the geographic context. Allopatric populations of *B. kinseyi* sampled from *Qv* exhibit significant habitat isolation with respect to the alternative host plant *Qg* (mean host preference and 95% CI: 0.629, 0.581– 0.676). Such habitat isolation in allopatric species might evolve when there is a fitness cost for searching for hosts (e.g., predation risk or highly variable environments) and selection increases the host search efficiency or reduces risk via increased host fidelity (Egan and Funk 2006). Similarly, the significantly higher estimate of sexual isolation among allopatric species pairs with different host associations (*B. kinseyi* × *B. fossoria*) compared to allopatric species with the same host associations (*B. kinseyi* × *B. treatae*) might arise as a by-product of phenotypic divergence (e.g., body size or abdomen size) due to local adaptation to different host plants (Egan et al. 2012a, Roush et al., in review).

Lastly, patterns of reproductive character displacement (RCD), where both habitat isolation and sexual isolation are higher among sympatric species with different host plant associations compared to allopatric species with different host plant associations, contributes a large proportion of total habitat isolation (46%) and sexual isolation (60%) between sympatric species (Fig. 5). Thus, RCD is likely to play a critical role in completing RI between *B. treatae* and *B. fossoria*. There are three notable mechanisms that might promote RCD: the Templeton effect, reinforcement, and reproductive interference (Templeton 1981; Noor 1999; Hollander et al. 2018). The Templeton effect, also known as differential fusion, posits that isolated populations experiencing strong prezygotic reproductive isolation persist in sympatry whereas weakly isolated populations fuse (Templeton 1981). Thus, the degree of prezygotic RI observed among sympatric species pairs represents a non-random sample of the range of RI between all species that have come into secondary contact. However, the Templeton effect is unlikely to be the cause of RCD in the *Belonocnema* system as it predicts elevated postzygotic reproductive isolation among sympatric species, which would prevent fusion of divergent species during secondary contact (Noor 1999). To date, we have no support for this pattern among *Belonocnema* species as the degree of hybrid inviability between sympatric *B. fossoria* and allopatric *B. kinseyi* is similar to the degree of hybrid inviability between sympatric species *B. fossoria* and *B. treatae* (Zhang et al. 2021a).

We argue that in the *Belonocnema* species complex that reproductive interference is a likely process that could promote RCD. In contrast to the process of reinforcement, which often occurs with ongoing gene flow between species, reproductive interference posits that the cost of migration, despite no current gene flow, could promote the elevation of prezygotic isolation between the two sympatric species (Butlin 1987; Hollander et al. 2018). Consistent with the prediction of reproductive interference, we found evidence of migration in a recent population genomic study, but no sign of gene flow between *B. treatae* and *B. fossoria* (Driscoe et al. 2019). And, we found that there was a cost of migration and mating in the form of immigrant inviability and hybrid inviaiblity (Zhang et al. 2017, 2021a). Moreover, given that the genetic divergence between sympatric species pairs is lower than allopatric species pairs, patterns of RCD indicate the strong effect of reproductive interference in overcoming the potential effect of larger genetic divergence in promoting reproductive isolation. What’s more interesting is that we observed that there is asymmetry in the direction of habitat isolation and sexual isolation between *B. treatae* and *B. fossoria* (Fig. 3A, Fig. S2), and that the direction of asymmetry is consistent with the direction of asymmetry in migration rate among these species (Nosil et al. 2003). More specifically, previous research has demonstrated lower migration of *B. fossoria* from *Qg* to the sympatric alterative host plant *Qv,* in comparison to the migration of *B. treatae* from *Qv* to the alternative host plant *Qg* (Driscoe et al. 2019; Zhang et al. 2021c). Correspondingly, the lineage that emigrates more (here, *B. treatae*) is predicted to evolve stronger habitat isolation due to the lower fitness of immigrants in the alternative habitat (Fig. 3A), while the lineage that receives more immigrants (here, *B. fossoria*) is predicted to exhibit stronger sexual isolation due to the lower fitness of hybrids (Fig. S2). Despite asymmetric migration and asymmetric RI being commonly found across different study systems (Bolnick et al. 2008; Lowry et al. 2008a; Oswald et al. 2017) and the understanding that migration rate is important with respect to reinforcement (Servedio and Noor 2003), few studies have investigated the consistency of the relationship between asymmetry in migration and asymmetry in promoting RI. Our study constitutes an example where the pattern of asymmetric RCD is predicted by the observed asymmetric migration rate (Yukilevich 2012; Suni and Hopkins 2018).

The results of the present study, when combined with previous studies of the *Belonocnema* wasp–live oak system, provide a case study to assess the relative contributions of pre-zygotic and post-zygotic barriers to total RI, which is a critical question in the biology of speciation (Coyne and Orr 2004; Nosil 2012; Coughlan and Matute 2020). Among sympatric *Belonocnema* species, multiple prezygotic barriers collectively generate near complete RI (total RI = 0.97, Egan et al. 2012 a,b, Hood et al. 2019, Zhang et al 2021a), whereas the strength of postzygotic barriers (hybrid inviability) is incomplete in comparison (Zhang et al. 2021a). While many studies have shown that prezygotic barriers can evolve faster than postzygotic barriers and therefore play a more important role in the early stage of speciation (Orr and Coyne 1989; Coyne and Orr 1997; Mendelson 2003; Rosser et al. 2019; Matute and Cooper 2021), our study provides an example where prezygotic barriers continue to be critical at the late stage of speciation (Jiménez-López et al. 2023). Moreover, a common perception is that genetic divergence plays a critical role in generating postzygotic isolation via genetic incompatibility (Orr and Coyne 1989; Coyne and Orr 2004; Coughlan and Matute 2020; Matute and Cooper 2021), while ecological selection contributes more to prezygotic isolation (Funk et al. 2006; Nosil 2012; Matute and Cooper 2021). In this regard, evidence found in the *Belonocnema*–live oak system challenges the traditional consensus on the role of ecology vs. genetic divergence in the evolution of pre vs. post reproductive isolation. First, several prezygotic barriers are the direct result of divergent selection, including reduced immigrant viability and fecundity and temporal isolation, which combine to contribute a large proportion of RI (0.87, Hood et al. 2019). Second, the postzygotic barrier ‘hybrid inviability’ between sympatric *B. treatae* and *B. fossoria* is subtle and varies among individual host plants, however, there is no hybrid inviability detected between allopatric *B. treatae* and *B. kinseyi* that share the same host. Together, these results strongly suggest that ecologically based selection, not genetic divergence, is the underlying mechanism of context-dependent hybrid inviability (Zhang et al. 2021a). However, mechanisms in promoting allopatric divergence such as genetic drift and non-host-associated divergence might play a much more important role in the two prezygotic reproductive barriers habitat isolation and sexual isolation than divergent ecology (Fig. 5). Lastly, as suggested by this study, geography plays an important role as an additional source of selection in sympatry and contributes significantly to both increased habitat isolation and sexual isolation (Fig. 5). Most studies on reinforcement focus on selection against hybrids resulting from genetic incompatibilities (Noor 1995; Jaenike et al. 2006; Hopkins and Rausher 2012; Barton 2020). In contrast, our finding provides a potential case where selection against hybrids and immigrants due to ecological adaptation to different host plants promotes the evolution of habitat isolation and sexual isolation. The finding that both ecology and geography play critical roles in completing speciation among *Belonocnema* species mirrors the few studies directly exploring this topic (e.g., stick insects: Nosil 2007; Guppies: Schwartz et al. 2010). However, more studies are needed to establish the general importance of these two evolutionary mechanisms in the process of speciation.

## Supporting information

Suppplement

## Author contributions

L.Z., and S.P.E. designed the study and all authors collected data. L.Z. analyzed the data and wrote the first draft of the manuscript. All authors edited subsequent drafts and approved the final version.

## Acknowledgements

We thank the staff and volunteers at Archbold Biological Station, especially Mark Deyrup and Hilary Swain; for field assistance in Florida; Robert Busbee, Amanda Driscoe, and Chelsea Smith for plant husbandry; Gaston Del Pino, Sean Liu, and Elaine Hu for data collection. The authors acknowledge Lake Lizzie Conservation Area in Osceola County and South Florida Water Management District at Kissimmee River for access to experimental sites. Support was provided to SPE by Rice University, a Rosemary Grant Award from the Society for the Study of Evolution (LZ), student research awards from the Society for Integrative and Comparative Biology, and the American Association of Naturalist (LZ).

## Notes

### Competing Interest Statement

The authors have declared no competing interest.

### Summary of Updates

Conceptual framework, all figures, and results

