## Supplementary material for "The role of ecology and geography in the evolution of habitat isolation and sexual isolation among sister species of host-plant-specific insects": Suppplement

**Supporting information**

**Table S1.** Details of all collection sites for *Belonocnema* wasps used in this study: site names (with abbreviations used in Table S3 and all Figures), U.S. state, host plant (*Qg* – *Quercus geminata*; *Qv* – *Quercus virginiana*); geographic context (sympatry – geographic overlap of host plants, allopatry – no geographic overlap of host plants, see map Figure 1); latitude and longitude.

| **Species** | **Site** | **State** | **Host plant** | **Geography** | **Latitude** | **Longitude** |
| --- | --- | --- | --- | --- | --- | --- |
| *B. fossoria* | Archbold Biological Station (ABS) | FL | *Qg* | sympatry | 27.1846 | -81.3521 |
| *B. fossoria* | Dickinson State Park (DCK) | FL | *Qg* | sympatry | 27.0261 | -80.1091 |
| *B. fossoria* | Lake Lizzie (LL) | FL | *Qg* | sympatry | 28.2276 | -81.1788 |
| *B. treatae* | Alva (Alva) | FL | *Qv* | sympatry | 26.7092 | -81.6154 |
| *B. treatae* | Cross City (CC) | FL | *Qv* | sympatry | 29.6192 | -83.0988 |
| *B. treatae* | Dauphin Island (DI) | AL | *Qv* | sympatry | 30.2516 | -88.0920 |
| *B. treatae* | Jekyll Island (JI) | FL | *Qv* | sympatry | 31.0175 | -81.4297 |
| *B. treatae* | Kissimmee river (KRE) | FL | *Qv* | sympatry | 27.3780 | -81.0968 |
| *B. treatae* | Okeechobee (Okee) | FL | *Qv* | sympatry | 27.2440 | -80.8168 |
| *B. kinseyi* | Golden Meadow (GM) | LS | *Qv* | allopatry | 29.3939 | -90.2729 |
| *B. kinseyi* | Picayune (PY) | MS | *Qv* | allopatry | 30.5255 | -89.6795 |
| *B. kinseyi* | Rice University (Rice) | TX | *Qv* | allopatry | 29.7174 | -95.4018 |

**Table S2.** Number of females tested per site to measure habitat isolation. Only females located on a host plant at least once during the 30 minute observation period were retained in the analysis of host preference.

| **Species** | **Site** | **# females tested in host preference trials** | **# females retained in host preference analysis** |
| --- | --- | --- | --- |
| *B. fossoria* | Archbold Biological Station (ABS) | 42 | 31 |
| *B. fossoria* | Dickinson State Park (DCK) | 46 | 40 |
| *B. fossoria* | Lake Lizzie (LL) | 42 | 38 |
| *B. treatae* | Cross City (CC) | 99 | 66 |
| *B. treatae* | Kissimmee river (KRE) | 27 | 14 |
| *B. kinseyi* | Golden Meadow (GM) | 82 | 62 |
| *B. kinseyi* | Picayune (PY) | 36 | 29 |
| *B. kinseyi* | Rice University (Rice) | 183 | 146 |
|  | **Total** | 557 | 426 |

**Table S3**. Number of males and females tested per site to measure sexual isolation in this study.

|  |  | ***B. fossoria* (**♀**)** | | |  | | ***B. treatae* (**♀**)** | | | | | |  | | ***B. kinseyi* (**♀**)** | | | |  | |
| --- | --- | --- | --- | --- | --- | --- | --- | --- | --- | --- | --- | --- | --- | --- | --- | --- | --- | --- | --- | --- |
|  | **Site** | **ABS** | **DCK** | **LL** | |  | | **Alva** | **DI** | **JI** | **KRE** | **Okee** | |  | | **GM** | **PY** | **Rice** | | **Total** |
| ***B. fossoria*** | **ABS** | 21 | 20 | 29 | |  | | 2 | 3 | 17 | 9 | 0 | |  | | 5 | 7 | 31 | | 144 |
| **(**♂**)** | **DCK** | 24 | 23 | 16 | |  | | 2 | 4 | 16 | 10 | 2 | |  | | 9 | 10 | 24 | | 140 |
|  | **LL** | 7 | 7 | 69 | |  | | 2 | 1 | 13 | 13 | 2 | |  | | 7 | 3 | 47 | | 171 |
| ***B. treatae*** | **Alva** | 3 | 13 | 5 | |  | | 7 | 0 | 0 | 2 | 3 | |  | | 0 | 0 | 10 | | 43 |
| **(**♂**)** | **JI** | 6 | 5 | 5 | |  | | 0 | 1 | 0 | 2 | 0 | |  | | 3 | 0 | 0 | | 22 |
|  | **KRE** | 8 | 9 | 7 | |  | | 1 | 1 | 3 | 15 | 0 | |  | | 0 | 0 | 19 | | 63 |
|  | **Okee** | 1 | 5 | 2 | |  | | 2 | 0 | 0 | 0 | 10 | |  | | 0 | 0 | 5 | | 25 |
| ***B. kinseyi*** | **GM** | 26 | 22 | 38 | |  | | 0 | 0 | 13 | 3 | 0 | |  | | 0 | 10 | 8 | | 120 |
| **(**♂**)** | **PY** | 12 | 17 | 21 | |  | | 0 | 0 | 14 | 2 | 0 | |  | | 5 | 0 | 8 | | 79 |
|  | **Rice** | 21 | 34 | 87 | |  | | 5 | 0 | 2 | 30 | 13 | |  | | 0 | 0 | 124 | | 316 |
|  | **Total** | 129 | 155 | 279 | |  | | 21 | 10 | 78 | 86 | 30 | |  | | 29 | 30 | 276 | |  |

**Table S4.** Results from General Linear Mixed Model (GLMM) analysis of host preference between different pairs of wasp species; (*df* = degrees of freedom). The term Method in the model is the effect of different testing environments (cup vs. Petri dish) in which the preference trials were conducted, while the term species indicates differences of host preference across different wasp species. Host preference differ significantly among different testing methods and wasp species, but not among interaction term.

| **GLMM host preference comparison** | | | |
| --- | --- | --- | --- |
|  | *χ*^2^ | *df* | *P*-value |
| **(Intercept)** | 6.637 | 1 | **0.010** |
| **Method** | 7.376 | 1 | **0.007** |
| **Species** | 6.330 | 2 | **0.042** |
| **Method:Species** | 2.065 | 2 | 0.356 |

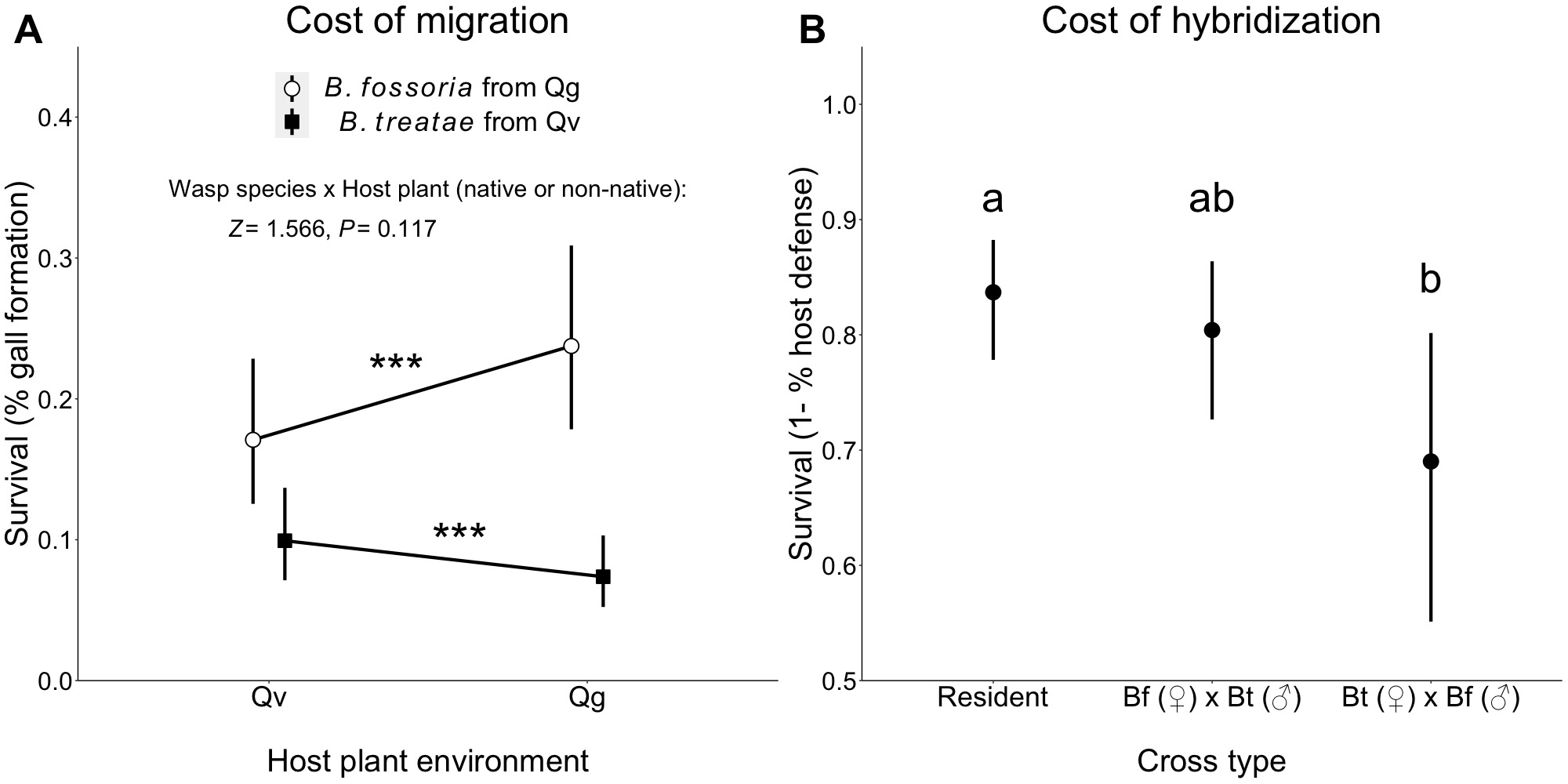

**Figure S1.** (A) Cost of migration for *B. fossoria* and *B. treatae* to alternative sympatric host plants as measured by the percent of oviposition events that form galls when females oviposit on their natal host plant compared to the alternative host plant (modified from Zhang et al. 2021a). Shown are least square means of % gall formation with 95 % CI. Triple asterisks indicate significant costs of migration for both species (*P* < 0.001). (B**)** Cost of hybridization as measured by probability of survival of individual eggs as a function of cross type (modified from Zhang et al. 2021b). Survival is indexed as the ability of developing larvae to evade the host plant’s hypersensitive immune response. The resident cross involves matings of conspecific males and females followed by oviposition into the natal host plants. All values compare the least squared mean with 95% CI. Means within panel labeled with different letters differ (*P* < 0.05).

**Methods and Results for Table S5, S6, and Figure S2**

**Testing for asymmetric sexual isolation among sympatric lineages**

Asymmetry in sexual isolation was tested by comparing the strength of mate preference between sympatric *B. treatae* and *B. fossoria* populations via GLMM in which male mate preference and female mate preference were evaluated separately. In assessing mating behavior in addition to copulation behavior, we also recorded two other courtship and mating-related interactions at each of the 15 observation intervals: whether the male was engaged in wing buzzing (or fanning), an important male courtship display (Villagra et al. 2011; Bredlau and Kester 2019), and whether the male mounted the female, a behavior that precedes copulation. Male preference was estimated by evaluating whether each individual displayed the wing buzz courtship behavior in one or more of the 15 two-minute intervals. Female preference was based on whether or not a female allowed the male to mount or copulate given that male courtship (wing buzzing) occurred during the 30-minute observation period (Egan et al. 2012*a*).

For each model, mate preference (Bernoulli trials: 0 or 1) was treated as the response variable with a binomial distribution, while the independent variables collection year, gall wasp species, species pair (conspecific or heterospecific), and the interaction term gall wasp species × species pair were included as fixed effects with collection site treated as a random effect. A significant interaction between species and species pair (conspecific versus heterospecific) is interpreted as evidence of asymmetric sexual isolation.

**Table S5.** Results of General Linear Mixed Model (GLMM) analysis of male mate preference among sympatric wasp species *B. treatae* and *B. fossoria*. The term “Male species” in the model indicates the species of male in each mating trial (*B. treatae* vs. *B. fossoria*) while the term “Species pair” indicates whether the male and female used in each trial were conspecifics or heterospecifics. For male mate preference, statistically significant differences were found across different collect year, male species. More importantly, the degree of mate preference difference between conspecifics and heterospecific individuals differ significantly across different male species.

|  | **χ^2^** | ***Df*** | ***P*-value** |
| --- | --- | --- | --- |
| **Intercept** | 5.537 | 1 | **0.019** |
| **Year** | 8.070 | 2 | **0.018** |
| **Male species** | 7.957 | 1 | **0.005** |
| **Species pair** | 0.592 | 1 | 0.442 |
| **Male species x Species pair** | 8.852 | 1 | **0.003** |

**Table S6.** Results GLMM for female mate preference among sympatric wasp species *B. treatae* and *B. fossoria*. Female species in the model indicates the species of female in each mating trial (*B. treatae* vs. *B. fossoria*) while “Species pair” indicates whether the male and female used in each trial were conspecifics or heterospecifics. For female mate preference, no statistically significant differences were found across different collect year, female species, or conspecifics vs. heterospecifics.

|  | **χ^2^** | ***Df*** | ***P*-value** |
| --- | --- | --- | --- |
| **Intercept** | 0.815 | 1 | 0.367 |
| **Year** | 3.625 | 2 | 0.163 |
| **Female species** | 0.407 | 1 | 0.523 |
| **Species pair** | 0.202 | 1 | 0.653 |
| **Female species × Species pair** | 0.002 | 1 | 0.960 |

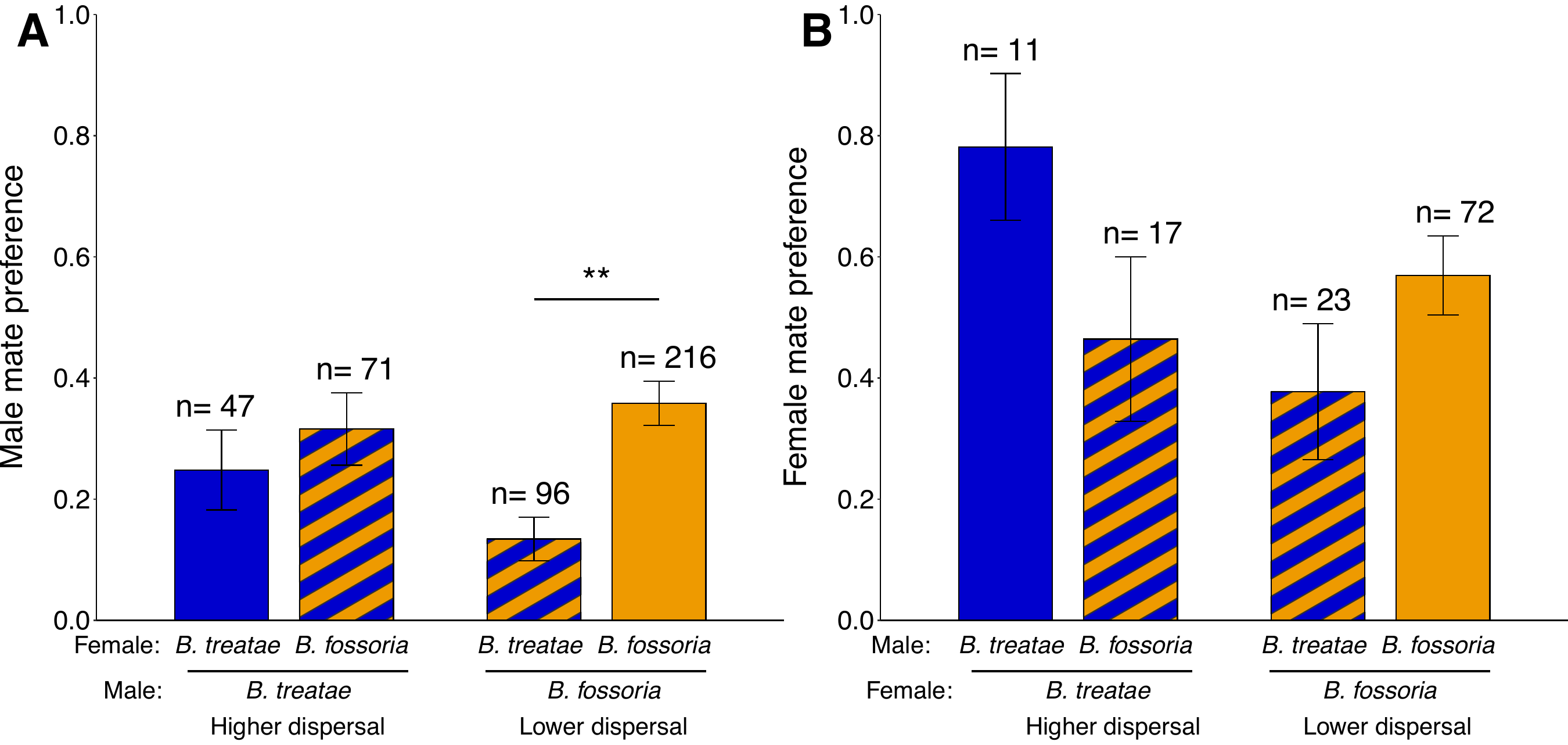

**Figure S2.** Strength of sexual isolation (mean ± SE) of males (A) and females (B) between sympatric lineages of *B. treatae* and *B. fossoria*. Solid-colored bars represent conspecific matings, and cross-hatched bars represent heterospecific matings. Asterisks indicate significant statistical differences of male mate preference between the same and different species pairs (*P* < 0.01). Asymmetrical male mate preference between *B. treatae* and *B. fossoria* is indicated by the significant male species × species pair interaction term in the GLMM (*P* = 0.0029), but no asymmetrical female mate preference was found (*P* = 0.960).
